# Circular DNA interacts with Cathelicidin LL37 to efficiently activate plasmacytoid dendritic cells through TLR9

**DOI:** 10.64898/2026.10.15.759370

**Authors:** Amara Ejikemeuwa, Jia He, Yuangao Wang, Guangming Li, Weirong Yuan, Wuyuan Lu, Yi Zhang, Lishan Su

## Abstract

Circulating cell-free DNA in inflamed tissues contains different forms of DNA, including extrachromosomal circular DNA (eccDNA). The role of DNA topology in innate immune sensing remains poorly understood. Here we show that circular DNA is a potent ligand for human plasmacytoid dendritic cells (pDCs) and that the cathelicidin host defense peptide LL37 selectively enhances circular DNA-induced activation. Across eccDNA, minicircle and synthetic circular DNA species, LL37 strikingly amplified type I interferon (IFN-I) and pro-inflammatory cytokine production, whereas linearized counterparts of identical sequence remained inactive. LL37-circular DNA complexes signaled through endosomal TLR9, as activation was abrogated by chloroquine and by the TLR9 antagonist IRS869. Mechanistically, LL37 promoted preferential retention of circular DNA in EEA1-positive early endosomes, and surface plasmon resonance revealed enhanced TLR9 binding to circular versus linear DNA, without differential LL37 binding. DNA topology rather than sequence, therefore, determines the immunostimulatory potential of cell-free DNA in pDCs. These findings define a mechanism by which LL37 and circular DNA jointly drive TLR9-dependent IFN-I responses, with implications for autoimmunity, cancer and nucleic acid therapeutics.

## Introduction

Plasmacytoid dendritic cells (pDCs), the body’s professional interferon-producing cells, respond to microbial encounter by rapidly producing robust amounts of type I interferon (IFN-I). This response is driven predominantly by the endosomal Toll-like receptors (TLRs) 7 and 9, which detect microbial RNA and DNA, respectively [1–3]. Although pDCs constitute only 0.2-0.8% of mononuclear cells in human peripheral blood and lymphoid tissues, chronic pDC activation and sustained IFN-α production contribute deleteriously to the pathogenesis of chronic viral infection, autoimmune diseases such as systemic lupus erythematosus (SLE) and psoriasis, and cancer [2–5] Tumor-associated pDCs are present in multiple cancers and are often linked to poor prognosis [2], as their recruitment supports the induction of regulatory T cells and an immunosuppressive tumor microenvironment [3].

Cell-free DNA (cfDNA) comprises extracellular DNA fragments released through diverse mechanisms, including multiple forms of tumor cell death and active secretion via extracellular vesicles [6]. Elevated cfDNA levels are observed in numerous diseases relative to healthy individuals, making cfDNA a promising biomarker for inflammatory disorders, including cancer [7, 8]. Beyond serving as a biomarker, cfDNA participates directly in shaping local immune responses. In persistently inflamed environments, recruited neutrophils can undergo NETosis, a specialized cell death program that releases extracellular traps composed of DNA, histones and antimicrobial peptides such as LL37 [9, 10]. LL37 is functionally promiscuous, exerting beneficial roles in combating bacteria, fungi and viruses, as well as detrimental effects that contribute to autoimmune disease and tumor progression [11, 12]. In autoimmune diseases such as SLE and psoriasis, LL37 forms complexes with cfDNA and potently activates pDCs via TLR9, driving chronic IFN-I production [13, 14]. Prior work on LL37-cfDNA-mediated pDC activation, however, has examined only linear DNA, even though cfDNA also contains extrachromosomal circular DNA (eccDNA) and mitochondria DNA.

It is reported that eccDNA is generated through DNA damage and repair pathways, hyper-transcription and DNA ligation after apoptotic fragmentation. It has attracted growing attention owing to its high prevalence in cancer, where it is associated with tumorigenesis and drug resistance, depending on the sequences it contains [15–17]. eccDNA is also found in other diseases, including cardiovascular disorders, inflammatory bowel disease and neurological disorders [18, 19]. When transfected by liposomes, eccDNA is a potent immunostimulant in bone marrow-derived dendritic cells and macrophages, whereas linearized eccDNA shows minimal activity [20]. Little is known about LL37 interactions with circular DNA, although atomic force microscopy suggests that LL37 can promote aggregation and coiling of plasmid DNA [13].

These findings led us to investigate the roles of LL37 and eccDNA in pDC activation. We observed that eccDNA was inherently more potent than linear DNA in activating pDCs and that LL37 further enhanced the activity of circular, but not linear, DNA. Our results indicate that LL37-circular DNA complexes activate pDCs through endosomal TLR9 signaling, likely due to LL37-mediated retention of circular DNA in the early endosome. In addition, circular DNA exhibited enhanced binding to TLR9 relative to linear DNA.

Together, these results provide insight into how DNA topology and host defense peptides shape the immunostimulatory potential of eccDNA, offering new perspectives on pDC activation in tumor and disease microenvironments.

## Results

### eccDNA is a potent immunostimulatory ligand for human pDCs, and LL37 selectively enhances its activity

Previous studies showed that, when delivered by liposomal transfection into bone marrow-derived dendritic cells and macrophages, eccDNA induces IL-6 and TNF-α [15, 20]. Whether eccDNA also stimulates pDCs was unclear. To address this, we first transfected eccDNA or linearized eccDNA (LinDNA) into total PBMCs, purified pDCs and pDC-depleted cells from two donors (Fig. S1A-C). Transfection of eccDNA, but not its linear counterpart, induced strong IFN-α responses—approximately 80-fold in purified pDCs of both donors (Fig. S1A) and approximately 150-fold in total PBMCs (Fig. S1B)—with no induction in the pDC-negative fraction (Fig. S1C).

Because transfection can bias the delivery of DNA ligands to nonphysiological subcellular compartments [21], we next examined eccDNA sensing under conditions that preserve endogenous uptake mechanisms. We directly incubated eccDNA or LinDNA with pDCs purified from independent donors (Fig. 1A). eccDNA alone, but not LinDNA, significantly induced IFN-I production (Fig. 1B-D). Strikingly, when LL37 was premixed with the DNA before incubation, it markedly enhanced eccDNA-mediated, but not LinDNA-mediated, IFN-I production (Fig. 1B-D). Consistent with this selectivity, LL37 enhanced eccDNA-mediated IL-6 (Fig. 2A, B) and TNF-α (Fig. 2C, D) production in purified pDCs. Together, these data demonstrate that eccDNA is a potent immunostimulatory ligand for pDCs and that LL37 selectively enhances circular DNA-mediated type I interferon and pro-inflammatory cytokine production.

**Figure 1.**
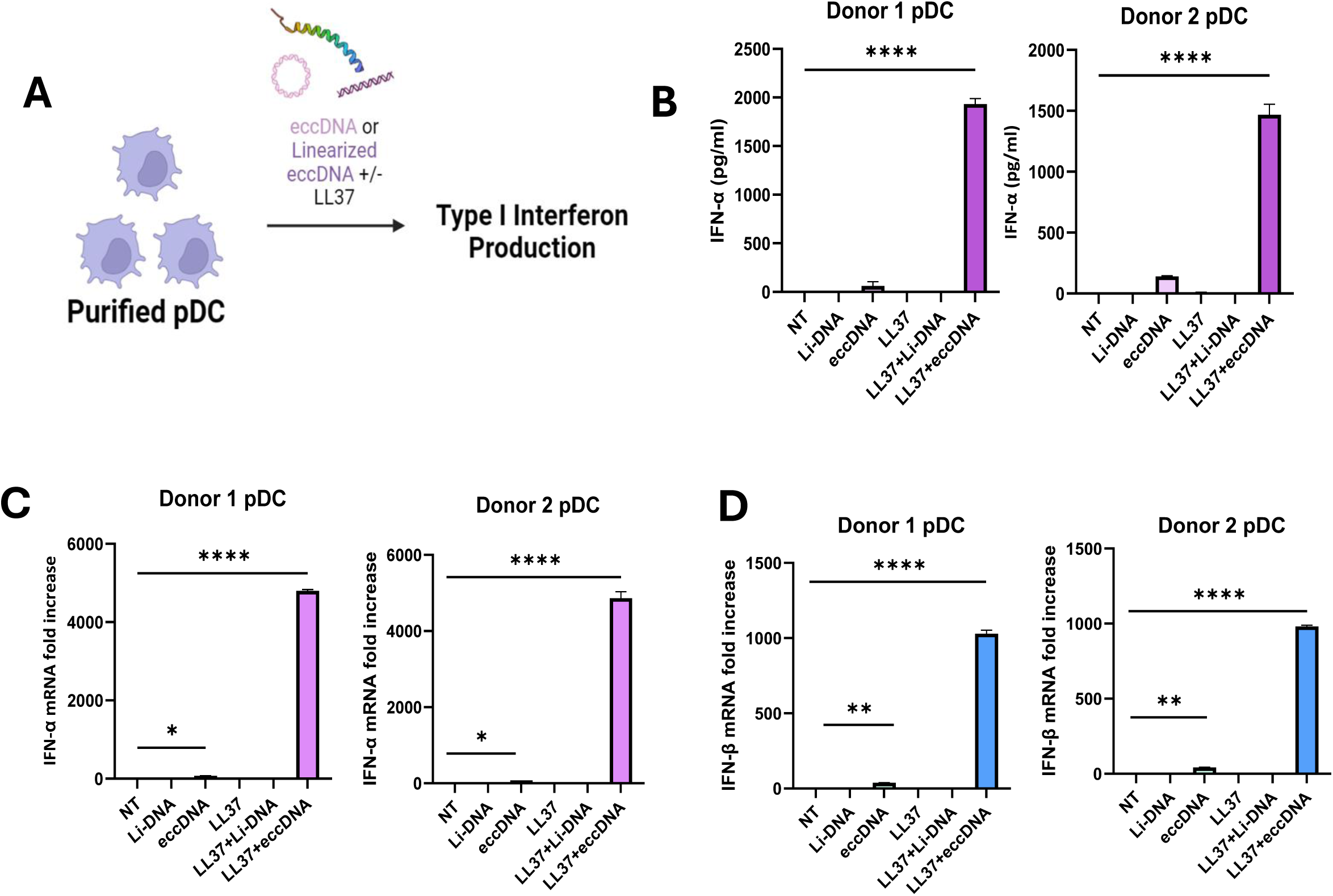
LL37 enhances eccDNA-mediated pDC activation. LL37 enhances eccDNA-mediated IFN-I production in pDCs. (A) pDCs were purified from two donor PBMC samples and incubated with eccDNA or linearized eccDNA (LinDNA) at 200 ng/mL with and without LL37 (2 µM) for 16 hours. (B–D) eccDNA, and not LinDNA, induced IFN-I production. IFN-α (B) protein was quantified by ELISA; mRNA expression of IFN-α (B) and IFN-β (C) was quantified by RT-PCR and normalized to GAPDH. Data were performed in duplicate with mean values ± SEM. *P < 0.05, **P < 0.01, ***P < 0.001 and ****P < 0.0001 by one-way ANOVA.

**Figure 2.**
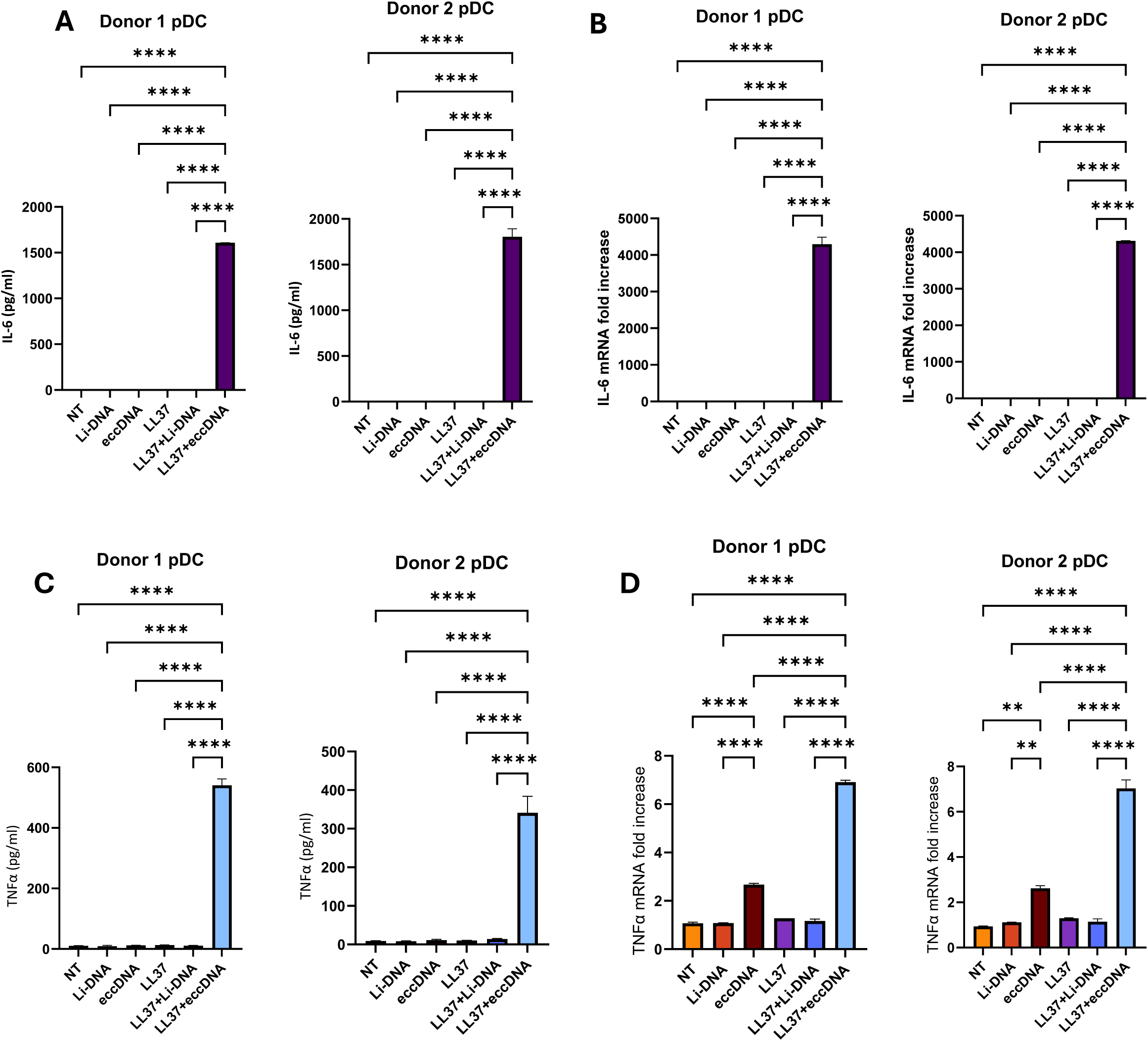
LL37 enhances eccDNA-mediated pDC pro-inflammatory cytokine production. LL37 enhances eccDNA-mediated IL-6 (A, B) and TNF-α (C, D) production in pDCs. pDCs were purified from two donor PBMC samples and incubated with eccDNA or linearized eccDNA (LinDNA) at 200 ng/mL with and without LL37 for 16 hours. IL-6 protein (A) and mRNA (B) quantification; TNF-α protein (C) and mRNA (D) quantification. Protein levels were quantified by ELISA; mRNA expression was quantified by RT-PCR and normalized to GAPDH. Data were performed in duplicate with mean values ± SEM. *P < 0.05, **P < 0.01, ***P < 0.001 and ****P < 0.0001 by one-way ANOVA.

### DNA topology, rather than sequence, determines LL37-enhanced pDC activation

To exclude the size and sequence heterogeneity inherent to eccDNA preparations,[22], we used circular DNA produced by a minicircle DNA system [23]. Purified pDCs from three donors were incubated with minicircle DNA (mc-DNA), linearized minicircle DNA (LinDNA), or each precomplexed with LL37 (Fig. 3A, B). As with eccDNA, LL37 significantly enhanced mc-DNA-mediated, but not linear mc-DNA-mediated, pDC activation in all donors (Fig. 3A, B).

**Figure 3.**
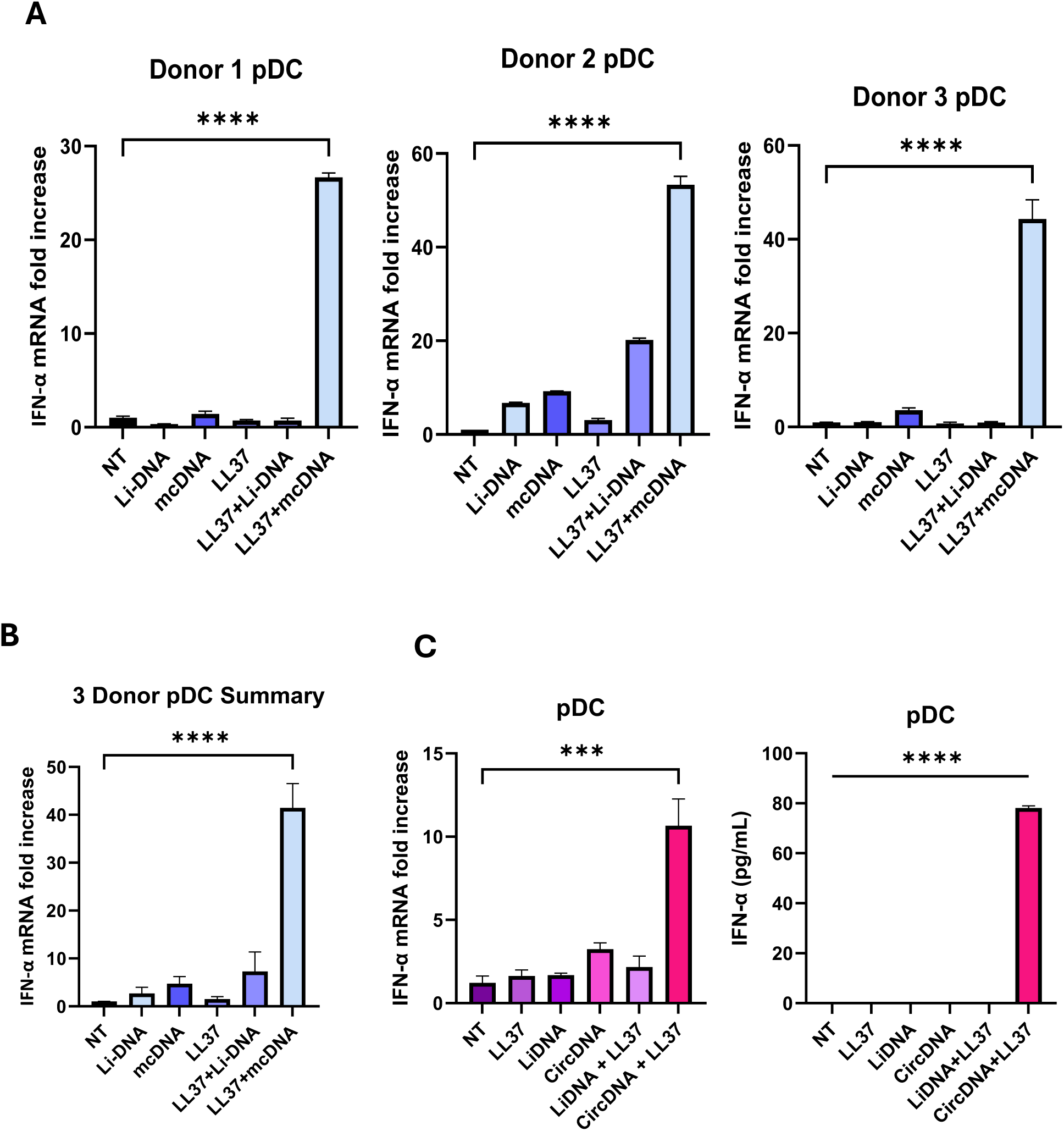
LL37 enhances multiple forms of circular DNA-mediated pDC activation. PBMCs from three human donors were used to purify pDCs. Minicircle DNA (mc-DNA) or linearized mc-DNA (LinDNA) was incubated (50 ng/mL) with purified pDCs for 16 hours. (A) LL37 enhances mc-DNA-mediated, but not LinDNA-mediated, IFN-α production in pDCs. (B) Three-donor pDC summary. (C) Synthetic minicircle ‘circDNA’ and its linear counterpart, ‘LinDNA’, ± LL37 were incubated with human pDCs. LL37 enhances circDNA-mediated IFN-I mRNA (left) and protein (right) expression in pDCs. Data were performed in duplicate with mean values ± SEM. *P < 0.05, **P < 0.01, ***P < 0.001 and ****P < 0.0001 by one-way ANOVA.

To establish further that circular topology, rather than DNA sequence, drives this response, we used a ligase-assisted minicircle accumulation (LAMA) method [24] to generate a 200-bp synthetic double-stranded circle, ‘circDNA’ (Supplementary Fig. 4). Stimulation of pDCs with LL37-circDNA complexes confirmed that LL37 enhances circular DNA-mediated pDC activation more efficiently than linear DNA in a sequence-independent manner (Fig. 3C). To facilitate mechanistic studies, we employed the pDC-like cell line Gen2.2, [25] and confirmed that LL37 enhanced activation by multiple forms of circular DNA—eccDNA (Supplementary Fig. 5A, B), minicircle DNAs (Supplementary Fig. 5C-E) and synthetic circDNA (Supplementary Fig. 5F, G)—while linear DNA had no effect. Collectively, these findings demonstrate that LL37 enhances circular, but not linear, DNA-mediated pDC activation regardless of size, biological source or sequence, in both primary pDCs and Gen2.2 cells, implicating circular topology itself as a determinant of immunostimulatory capacity in pDCs.

### LL37-circDNA sensing is dependent on endosomal TLR9

CpG oligonucleotides and linear DNA activate pDCs through endosomal TLR9. To determine whether LL37-circular DNA complexes use a similar pathway, we blocked endosomal acidification with chloroquine (CQ), which inhibits endosome maturation and signaling [26] (Fig. 4; Supplementary Fig. 6). CQ treatment markedly reduced LL37-circular DNA-mediated pDC activation for both eccDNA (Fig. 4A) and mc-DNA (Fig. 4B). To confirm the involvement of TLR9 itself, we inhibited TLR9 signaling with the antagonist IRS869 [27, 28]. IRS869 markedly reduced LL37-circDNA-mediated IFN-α production in PBMCs (Fig. 4C), and LL37-circDNA-induced IFN-α and IL-6 expression in Gen2.2 cells was significantly attenuated (Fig. 4D, E). These responses are therefore TLR9 dependent.

**Figure 4.**
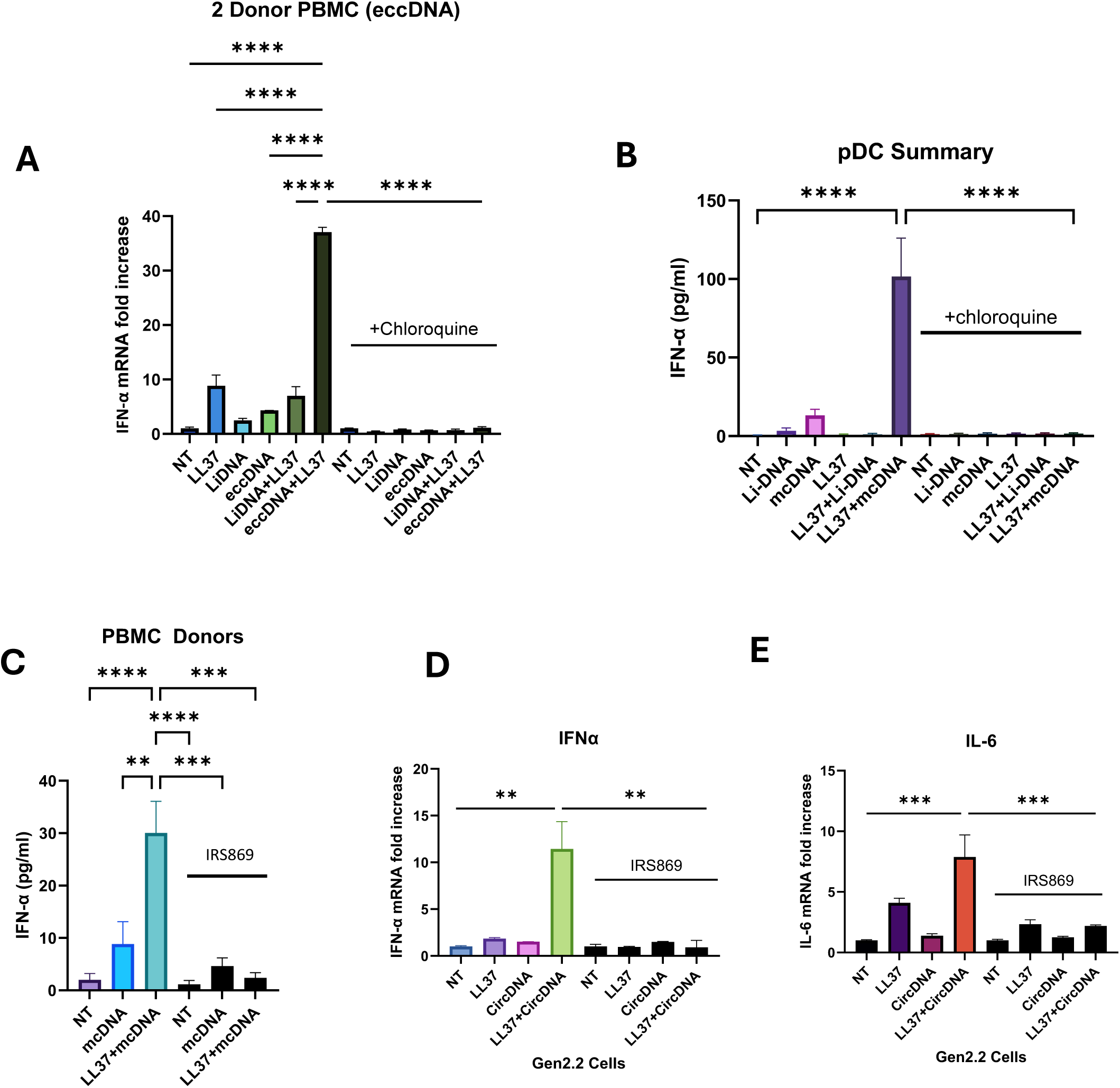
LL37-circDNA sensing is dependent on endosomal TLR9 stimulation. Purified PBMCs (A, C) or pDCs (B, D, E) were preincubated with the endosomal acidification inhibitor chloroquine (CQ; A, B) or the TLR9 antagonist IRS869 (C–E) for 1 hour, then washed and stimulated. CQ inhibits LL37–eccDNA-mediated pDC activation in 2 PBMC donors (A) and LL37–mc-DNA-mediated pDC activation in all 3 donors (B). IRS869 inhibits LL37–mcDNA-mediated PBMC activation (C) and LL37–circDNA-mediated Gen2.2 IFN-α (D) and IL-6 (E) production. IFN-α was quantified by ELISA and RT-PCR; RT-PCR was normalized to GAPDH. Data were performed in duplicate with mean values ± SEM. *P < 0.05, **P < 0.01, ***P < 0.001 by one-way ANOVA.

### LL37 leads to preferential retention of circDNA within the early endosomal compartment

LL37 modulates DNA trafficking within endosomes to affect TLR9 signaling [13, 29]. CpG-B, a late endosomal TLR9 agonist, adopts CpG-A-like behavior when complexed with LL37, shifting from NF-κB-mediated pro-inflammatory responses to an IFN-I-dominant program [13]. Given that LL37-circDNA induces both IFN-I and pro-inflammatory cytokines, we asked whether LL37 alters the endosomal trafficking of circular relative to linear DNA. Gen2.2 cells were incubated with 488-dUTP-labeled linear or circular DNA ± LL37 for 6 hours, followed by confocal analysis of colocalization with the early endosome marker EEA1 and the late endosome marker LAMP1 (Fig. 5A–D). In the presence of LL37, circDNA exhibited a significantly higher EEA1/LAMP1 ratio than circDNA alone or LinDNA ± LL37 (Fig. 5E). Consistently, LL37 induced a significantly greater increase in EEA1 colocalization for circDNA than for LinDNA (Fig. 5F), indicating preferential retention of circDNA within the early endosomal compartment—the compartment in which TLR9 engagement occurs.

**Figure 5.**
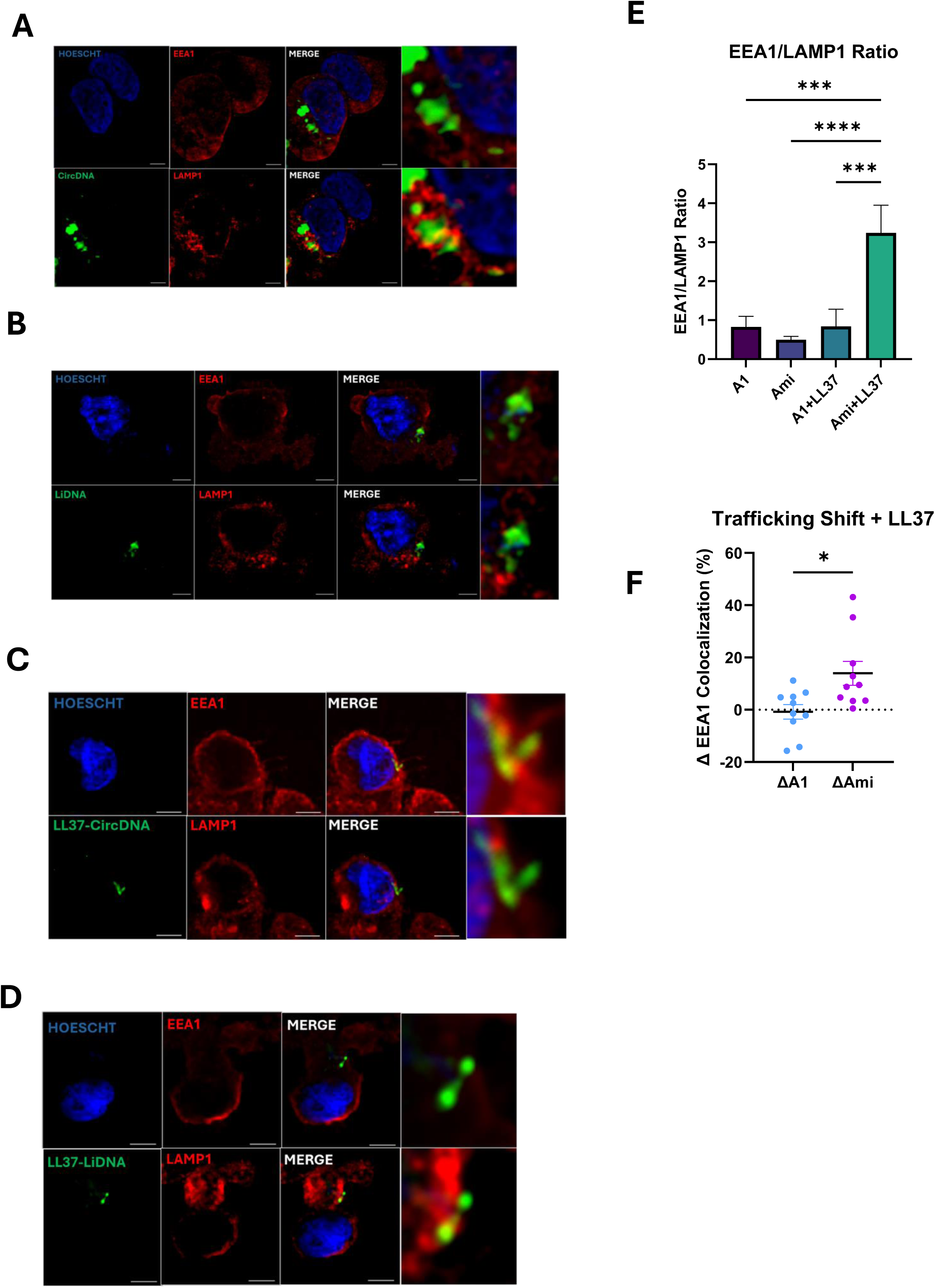
LL37 leads to preferential retention of circDNA within the early endosomal compartment. Gen2.2 cells were stimulated for 6 h with CF488-dUTP-labeled linear DNA (LinDNA) or circular DNA (circDNA) alone or precomplexed with LL37. Cells were stained for the early endosome marker EEA1 (Alexa Fluor 647, red), the late endosome marker LAMP1 (Alexa Fluor 594, red), and nuclei were counterstained with Hoechst (blue). (A–D) Representative 40× confocal images of LinDNA or circDNA ± LL37 showing DNA localization relative to EEA1 or LAMP1; enlarged regions of interest are shown in the right panels. (E) Quantification of the EEA1/LAMP1 colocalization ratio. (F) Change in EEA1 colocalization (ΔEEA1) following LL37 treatment for LinDNA and circDNA, demonstrating a greater LL37-induced increase in early endosomal localization of circDNA. Images were acquired using a Nikon W1 spinning-disk confocal microscope. Colocalization was quantified from 10 fields using Fiji following image processing in ZEN Blue. Data are presented as mean ± SEM; statistical significance was determined by one-way ANOVA. Line in images represents 5 µm distance.

### TLR9 exhibits enhanced binding to circular DNA compared with linear DNA

To assess whether DNA topology influences interactions with LL37 and TLR9, we evaluated binding by surface plasmon resonance (SPR) (Fig. 6). DNA was biotinylated by PCR incorporation to generate linear fragments, or further ligated to form circular DNA, and immobilized on SPR sensor chips. LL37 alone exhibited comparable binding to linear and circular DNA, with minimal differences in affinity (KD) and maximal binding response (Rmax) (Fig. 6A, B, E), and LL37-DNA complexes displayed similarly prolonged dissociation half-lives independent of DNA topology (Fig. 6E). In contrast, SPR analysis showed a trend toward higher affinity, increased Rmax and a longer dissociation half-life for circDNA compared with linear DNA at matched concentrations (Fig. 6C-E), consistent with a preferential interaction between TLR9 and circDNA. Together, these findings support a model in which LL37 enhances circDNA signaling by promoting its localization within the early endosomal compartment, where TLR9 engagement is likely to occur (Fig. 6F). Although LL37 does not differentially bind linear and circular DNA, circDNA itself trends toward stronger affinity and a longer dissociation half-life with TLR9 than linear DNA. These spatial and binding differences provide a mechanistic basis for the enhanced TLR9-dependent inflammatory responses induced by LL37-circDNA complexes.

**Figure 6.**
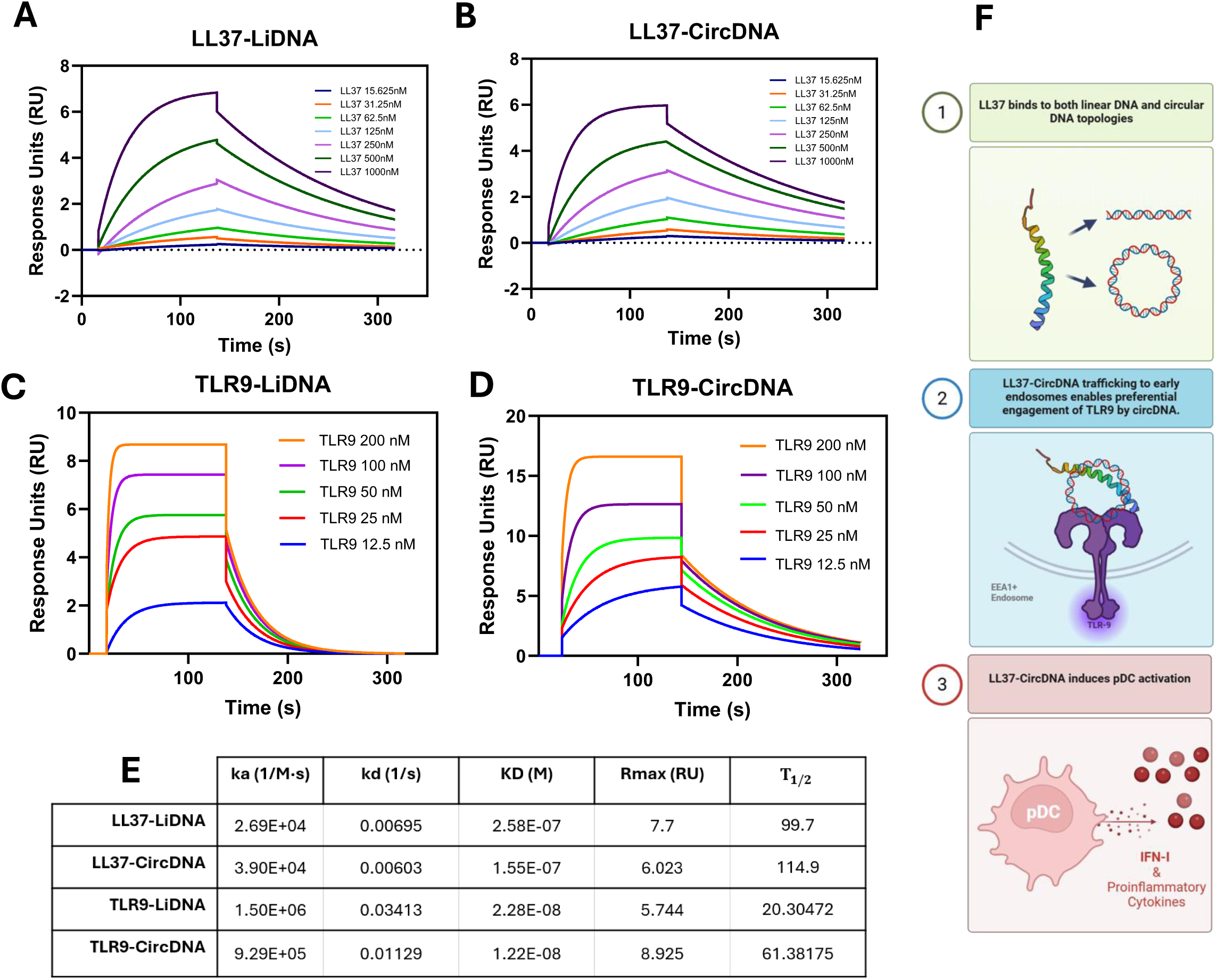
TLR9 exhibits enhanced binding properties with circDNA compared with linear DNA. Surface plasmon resonance (SPR) sensorgrams of (A) LL37–LinDNA, (B) LL37– circDNA, (C) TLR9–LinDNA, (D) TLR9–circDNA, (E) SPR summaries. (F) Schematic summarizing the proposed mechanism by which LL37 enhances circular DNA-mediated TLR9 activation. SPR data from the Biacore Core Facility; schematic made in BioRender.

## Discussion

pDCs walk a fine line between disease prevention and progression. In acute viral infection, they serve as powerful sentinels bridging innate and adaptive immunity through rapid IFN-I secretion, recruitment of immune effectors and T cell co-stimulation [1, 30]. When their IFN response is dysregulated or misdirected toward self-nucleic acids, however, pDCs promote chronic inflammation and autoimmunity [1–3, 13, 14, 25, 27, 30–37]. This duality is particularly relevant in conditions where extracellular nucleic acids accumulate, such as autoimmune disease, infection and cancer.

Neutrophils are constantly recruited and activated in diseased environments and can undergo NETosis, during which peptides such as LL37 are released [11] [12]. LL37 is a 4.5-kDa amphipathic α-helical peptide that binds linear DNA by forming a cationic backbone that coordinates multiple DNA strands into ordered structures [38]. Atomic force microscopy has further suggested that LL37 induces coiling and aggregation when bound to circular DNA, although this remains poorly understood [13]. Our findings provide functional insight into these structural observations, demonstrating that LL37 preferentially enhances circular, but not linear, DNA-mediated IFN-I production in pDCs.

In this study, we demonstrate that eccDNA significantly activated pDCs to produce type I interferons and pro-inflammatory cytokines, whereas linearized eccDNA did not, and that this activation is strikingly enhanced by LL37. LL37 enhanced pDC activation across multiple forms of circular DNA with unrelated sequences—eccDNA, minicircle DNA and synthetic circular DNA—but not their linear counterparts. Notably, these effects occur at low DNA concentrations (1–200 ng/mL), within the physiological range of cell-free DNA reported in disease contexts [39] [40]. By contrast, prior studies examining LL37 with linear DNA, or circular DNA alone, often used substantially higher DNA concentrations (10–50 µg/mL) [13, 14, 37], which may not reflect physiological conditions. As evidenced by sensitivity to chloroquine and the TLR9 antagonist IRS869, together with TLR9 binding assays and fluorescence microscopy, circDNA sensing is dependent on endosomal TLR9 signaling.

These results carry broad implications for cancer and autoimmune disease biology. eccDNA has emerged as a key player in tumorigenesis, arising from chromosomal breakage, replication stress, defective repair or apoptotic DNA fragmentation [19, 20, 41, 42]. These circular DNA elements can contain oncogenes or regulatory sequences that promote genetic heterogeneity, transcriptional plasticity and therapy resistance[16, 43]. Excessive extrachromosomal DNA and LL37 are also present in autoimmune diseases such as SLE, rheumatoid arthritis, type 2 diabetes mellitus and psoriasis [13, 14, 33–35, 38, 44–46]. LL37-circular DNA complexes in these disease contexts have yet to be investigated.

Beyond its genomic contribution to disease, eccDNA is increasingly recognized as an immunostimulant in macrophages and dendritic cells [15, 20]. Sequencing studies have shown that eccDNAs originate from all chromosomes and span a wide range of sizes, from approximately 188 bp to more than 15 kb [15, 20]. Importantly, our findings show that LL37 enhancement of pDC activation occurs across diverse circular DNA species (200 bp to >8 kb), sequences and biological sources, further supporting DNA topology— rather than sequence or length—as a major determinant of pDC immunogenicity.

Mechanistically, our data indicate that the enhanced immunostimulatory activity of LL37– circDNA complexes is not driven by preferential LL37 binding to circular versus linear DNA, whereas TLR9 showed preferential interaction with circDNA over LinDNA. In addition, LL37 promoted the localization of circDNA within the early endosomal compartment, where TLR9 activation occurs. This finding is consistent with previous studies demonstrating that LL37 can alter the intracellular trafficking and compartmentalization of nucleic acids [13, 34, 35]. Our results suggest that LL37 provides an additional layer of regulation by modifying the intracellular trafficking of eccDNA, thereby enhancing or redirecting its immunostimulatory activity.

Importantly, the consequences of LL37-mediated trafficking may be cell-context dependent, potentially promoting or restricting nucleic acid sensing depending on the trafficking pathways and innate immune sensors engaged within a given cell population. LL37 may therefore function not simply as an enhancer of eccDNA sensing, but as a modulator that helps determine the immunostimulatory outcome of eccDNA in different cellular contexts; how this function extends to other immune cell populations remains to be investigated. Independently, our binding studies support prior work showing preferential interaction of TLR9 with curved DNA compared with linear DNA [47, 48]. Together, these findings support a model in which LL37 regulates the intracellular trafficking of circDNA, whereas DNA topology determines the efficiency of TLR9 engagement.

From a translational perspective, our results raise important considerations for nucleic acid-based therapeutics, gene delivery systems and vaccine platforms. Many of these technologies rely on circular DNA vectors, including plasmids and minicircles. Our findings suggest that such agents may inadvertently trigger pDC activation and IFN-I responses, particularly in the presence of endogenous peptides such as LL37. Conversely, this pathway could be harnessed for therapeutic benefit: controlled delivery of peptide-circular DNA complexes may enhance dendritic cell activation and immune priming in vaccine or cancer immunotherapy settings. However, therapies that induce eccDNA formation, such as genotoxic agents or radiation, may also carry unintended immunostimulatory consequences if eccDNA is released and sensed by pDCs.

While this study defines a role for LL37 in enhancing circular DNA-mediated pDC activation, several questions remain. The downstream signaling mechanisms that distinguish IFN-I and pro-inflammatory outputs were not fully resolved. Whether LL37 directly alters TLR9 receptor organization or accessibility within endosomal compartments also remains to be determined. Although our data support endosomal TLR9 engagement, additional pathways may contribute, as circular DNA has been shown to activate cytosolic sensors such as STING in other immune cell types [20, 49, 50]. Furthermore, in vivo studies will be required to determine the physiological relevance and potential pathological consequences of this pathway.

In summary, we identify DNA topology as a critical determinant of innate immunogenicity in plasmacytoid dendritic cells. We propose a model in which LL37 promotes localization of circDNA to the early endosomal compartment, while circular DNA topology facilitates efficient TLR9 engagement, together resulting in enhanced type I interferon responses. These findings provide a mechanistic framework for understanding how endogenous circular DNA species, including eccDNA, contribute to immune activation across disease contexts and highlight opportunities to therapeutically modulate this pathway.

## Methods

### Cell culture

Human buffy coats were obtained from the Gulf Coast Regional Blood Center. Primary human peripheral blood mononuclear cells (PBMCs) were isolated from buffy coats by Ficoll-Paque gradient centrifugation. BDCA-4/Neuropilin Microbead Kit (Miltenyi Biotec) and Plasmacytoid dendritic cell II isolation bead kits (Miltenyi Biotec) were used to isolate human pDCs from PBMCs. The pDC-like Gen2.2 cell line [26] was grown in suspension on a monolayer of MS5 feeder cells and maintained in RPMI 1640 medium containing 10% FBS, 1% penicillin/streptomycin, 2 mM glutamine and 10 mM HEPES (GIBCO).

### Circular DNA preparation

eccDNA was generated as reported [51]. The minicircle DNA production method was described in [23]. The pre-mcDNA in the plasmid backbone can be amplified in E. coli. After recombinase induction, pre-mcDNA undergoes attP–attB recombination and produces two DNA circles; the circle containing the bacterial backbone is degraded by I-SceI, while mcDNA is maintained. mc-cccDNA was linearized using the Fast Digestion Kit (Thermo Scientific) protocol with the XhoI restriction enzyme. Synthetic small DNA circles (‘circDNA’) were prepared by the ligase-assisted minicircle accumulation (LAMA) procedure described in [20]. The 200bp-dsDNA sequence was randomly generated (https://faculty.ucr.edu/~mmaduro/random.htm). The isomers of single-strand templates (LinDNA: 1 and 2) and their amplification primer sets were synthesized by IDT. Products with a 5′-end phosphate were prepared with 2× Q5 DNA polymerase (NEB). 400 ng of each isomer were added to generate 50-µL HiFi Taq DNA ligase reaction mixtures, which were placed in thermocyclers using the following cycles: 95 °C for 3 min, 60 °C for 10 min and 37 °C for 5 min, for 10 cycles. Circularized products were recovered with a PCR Purification Kit (Qiagen), digested with Plasmid-Safe ATP-dependent DNase (Lucigen) and recovered again with a PCR Purification Kit.

### LL37–DNA stimulation

2 × 10^5 PBMCs, 2 × 10^4 purified pDCs or 2.5 × 10^4 Gen2.2 cells were seeded in 96-well round-bottom plates in complete RPMI (Thermo Fisher Scientific, Waltham, MA) supplemented with 10% fetal bovine serum (FBS; Thermo Fisher Scientific), 2 mM glutamine (Thermo Fisher Scientific), 10 U/mL penicillin and 10 µg/mL streptomycin (Thermo Fisher Scientific). LL37 was generated in-house at the Institute of Human Virology, University of Maryland [52]. LL37 (final concentration 2–4 µM) and linear or circular DNA (final concentration 1–200 ng/mL) were premixed to form complexes in PBS for 30 min before being added directly to cell cultures. Commercial LL37 (InvivoGen) was used to validate in-house LL37 (Supplementary Fig. 7). To inhibit endosome maturation or TLR9 activation, cells were preincubated with 50 µM chloroquine (MedChemExpress) or 5 µM IRS869 (InvivoGen), respectively, for 1 hour. Cells were then washed, resuspended in complete RPMI and stimulated overnight.

### Measurement of supernatant cytokine production

Supernatants of stimulated PBMC, pDC and Gen2.2 cultures were collected after overnight culture. Cytokine levels in the supernatant were determined using ELISA kits for human IFN-α (MABTECH), IL-6 and TNF-α (R&D Systems).

### Real-time PCR

Cellular RNA was isolated with RNeasy kits (Qiagen). cDNA was synthesized with SuperScript III, and qPCR was performed with SYBR Green Master Mix (Thermo Fisher Scientific). For real-time PCR detection of IFN-I expression, the primers were as follows: IFN-α—F: GCCTCGCCCTTTGCTTTACT, R: CTGTGGGTCTCAGGGAGATCA; IFN-β— F: GTGCCTGGACCATAGTCAGAGTGG, R: TGTCCAGTCCCAGAGGCACAGG; IL-6— F: CGGCCTTCCCTACTTCACAA, R: TTGCCATTGCACAACTCTTTTC; TNF-α—F: GTCCCCAAAGGGATGAGAAGT, R: TTTGCTACGACGTGGGCTAC. Expression was normalized to GAPDH—F: CATGAGAAGTATGACAACAGCCT, R: AGTCCTTCCACGATACCAAAGT.

### Immunofluorescence microscopy

To visualize the intracellular localization of DNA in pDCs, LinDNA was labeled with CF488-dUTP nucleotides (Biotium) via PCR and ligated via LAMA to generate labeled circDNA. Gen2.2 cells were stimulated with DNA alone or precomplexed with LL37 for 6 hours. Following stimulation, cells were washed extensively and resuspended in PBS. 15-mm circle coverslips were precoated with poly-L-lysine (MilliporeSigma) overnight and washed before spinning Gen2.2 cells down onto the coverslips. After attachment, cells were fixed in 4% paraformaldehyde (Thermo Fisher Scientific), permeabilized with 0.1% Triton X-100 (MilliporeSigma) and blocked in BSA-containing buffer. Cells were stained with rabbit anti-LAMP1 (Abcam) and mouse anti-EEA1 (Santa Cruz Biotechnology) followed by Alexa Fluor 647 and Alexa Fluor 594 fluorescent secondary antibodies, and nuclei were counterstained with Hoechst. For confocal microscopy, samples were imaged using a Nikon Spinning Disk Confocal (W1) equipped with appropriate laser lines and emission filters. Images were acquired using sequential excitation to minimize spectral overlap.

### Surface plasmon resonance

To assess LL37, TLR9 and DNA interactions, surface plasmon resonance (SPR) experiments were performed at the University of Maryland, Baltimore Biacore Core Facility. LinDNA was biotinylated through Biotin-11-dUTP incorporation during PCR (Thermo Scientific), and biotinylated circular DNA was generated thereafter via ligase-assisted minicircle accumulation (LAMA). Biotinylated DNA was immobilized onto streptavidin-coated sensor chips, and binding experiments were carried out by flowing analytes over the DNA-coated surface. LL37 or recombinant human TLR9 ectodomain (Creative Biomart) was injected at the indicated concentrations, and binding responses were recorded in real time. Sensorgrams were analyzed to determine binding kinetics and response profiles for linear and circular DNA substrates.

### Statistics and reproducibility

Experiments were independently repeated at least twice with similar results. Representative images are shown where indicated. Quantitative data, including ELISA and RT-qPCR measurements, are presented as mean ± SEM. Experiments were performed with duplicate measurements unless otherwise indicated. The number of independent human donors used for each experiment is indicated in the corresponding figure legend. Comparisons between two groups were performed using two-sided unpaired t-tests. For comparisons involving three or more groups, one-way ANOVA was performed followed by Tukey’s multiple comparisons test. Statistical significance was defined as P < 0.05. All statistical analyses were performed and graphs generated using GraphPad Prism version 10.

## Data availability

All data supporting the findings of this study are available within the article and its Supplementary Information, or from the corresponding author upon reasonable request. Source data files for Figs. 1–6 will be provided with the paper.

## Code availability

No custom code was used.

## Acknowledgements

We thank all the members of the Su laboratory for helpful discussions. We also acknowledge the University of Maryland School of Medicine Confocal Microscopy Core and the University of Maryland, Baltimore Biacore Core Facility for their assistance. We thank Dr. Yinghua Zhang for assistance with surface plasmon resonance experiments and data analysis. This work was supported in part by grants from National Institutes of Health grants CA228172 (to LS) and T32CA154274 (to Toni Antalis and Curt Civin).

## Author contributions

A.E. and J.H. designed and performed experiments, analyzed and interpreted the data, prepared the figures. A.E. wrote the manuscript. G.L. provided technical assistance with pDC isolation. W.Y. and W.L. generated LL37. Y.W. and Y.Z. generated and provided eccDNA and assisted with revisions. L.S. conceptualized the project, supervised the project, contributed to experimental design and data interpretation, revised the manuscript and acquired funding.

## Supplementary figure legends

**Supplementary Figure 1 | Transfection of eccDNA activates pDCs.**

eccDNA or linearized eccDNA (LinDNA) was transfected via Lipofectamine into PBMCs from two separate donors (A), pDCs purified from two donor PBMC samples (B), and pDC-negative PBMC fractions from two donors (C) for 16 hours. Transfection of eccDNA induces IFN-α in PBMCs (A) and pDCs (B). Transfection of eccDNA or LinDNA did not induce IFN-α in the pDC-negative PBMC fraction (C). dG:dC = transfection positive control (A–C). IFN-α was quantified by RT-PCR and normalized to GAPDH. Data were performed in duplicate with mean values ± SEM. *P < 0.05, **P < 0.01, ***P < 0.001 and ****P < 0.0001 by one-way ANOVA.

**Supplementary Figure 2 | pDC purification.**

pDC purification examples. pDCs were isolated from donor buffy coats via (A) BDCA-4/Neuropilin Microbead Kit (Miltenyi Biotec) and (B) Plasmacytoid dendritic cell II isolation bead kits (Miltenyi Biotec). Cells were stained with FITC or PE-Cy7-CD123 and PE or PB-CD4, and cell purity was verified by flow cytometry with analysis in FlowJo. Experiments proceeded with pDC purity of >90%.

**Supplementary Figure 3 | LL37–mc-DNA induces rapid IFN-I production in PBMCs.**

mcDNA ± LL37 was incubated with donor PBMCs for 16 hours. IFN-α (A) and IFN-β (B) mRNA expression was quantified by RT-qPCR and normalized to GAPDH. Data were performed in duplicate with mean values ± SEM. *P < 0.05, **P < 0.01, ***P < 0.001 and ****P < 0.0001 by one-way ANOVA.

**Supplementary Figure 4 | Ligase-assisted minicircle accumulation of circDNA.**

Generation of a 200-bp dsDNA synthetic minicircle from linear DNA (‘LinDNA’). Synthetic small DNA circles (‘circDNA’) were prepared by the ligase-assisted minicircle accumulation (LAMA) procedure described in^21^. The 200-bp dsDNA sequence was randomly generated (https://faculty.ucr.edu/~mmaduro/random.htm). The isomers of single-strand templates and their amplification primer sets were synthesized by IDT. Products with a 5′-end phosphate were prepared with 2× Q5 DNA polymerase (NEB). 400 ng of each isomer were added to generate 50-µL HiFi Taq DNA ligase reaction mixtures and placed in thermocyclers using the following cycles: 95 °C for 3 min, 60 °C for 10 min and 37 °C for 5 min, for 10 cycles. Circularized products were recovered with a PCR Purification Kit (Qiagen) and digested with Plasmid-Safe ATP-dependent DNase (Lucigen) before being recovered with a PCR Purification Kit. (A) Linear DNA and (B) circular DNA were titrated ± LL37 to identify conditions matching primary pDC IFN-I results. IFN was quantified by ELISA. Data represent experiments performed in duplicate with mean values ± SEM. *P < 0.05, **P < 0.01, ***P < 0.001 and ****P < 0.0001 by one-way ANOVA.

**Supplementary Figure 5 | LL37 enhances circular DNA-mediated Gen2.2 activation.**

Circular or linear DNA of different sequences was tested in Gen2.2 cells with and without LL37. LL37 enhanced circular eccDNA-mediated IFN-α (A) and IFN-β (B); minicircle HBV cccDNA-mediated IFN-α (C) and IFN-β (D); minicircle IL-4-mediated IFN-α (E); and synthetic minicircle ‘circDNA’-mediated IFN-I mRNA (F) and IFN-α protein expression (G). IFN-α, IFN-β and IL-6 expression was quantified by RT-PCR and normalized to GAPDH; IFN-α protein was quantified by ELISA. Data represent experiments performed in duplicate with mean values ± SEM. *P < 0.05, **P < 0.01, ***P < 0.001 and ****P < 0.0001 by one-way ANOVA.

**Supplementary Figure 6 | Chloroquine inhibits LL37–mcDNA-mediated pDC activation.**

Total PBMCs were isolated from three buffy coat donors and pDCs were positively selected and purified. (A) pDCs were preincubated with the endosomal acidification inhibitor chloroquine (CQ) for 1 hour, then washed and stimulated with mcDNA. (B) CQ inhibits LL37–mcDNA-mediated pDC activation in three donors. IFN-α was quantified by ELISA. Data were performed in duplicate with mean values ± SEM. *P < 0.05, **P < 0.01, ***P < 0.001 by one-way ANOVA.

**Supplementary Figure 7 | Confirmation of in-house versus commercial LL37.**

LL37 generated in^52^ was used for this study. Commercial LL37 (InvivoGen) was used to validate in-house LL37 (‘Lab’) with Gen2.2 cells (A). IFN was quantified by RT-PCR and normalized to GAPDH. Data represent experiments performed in duplicate with mean values ± SEM. *P < 0.05, **P < 0.01, ***P < 0.001 and ****P < 0.0001 by one-way ANOVA.

## References

1. Ye, Y., et al., Plasmacytoid dendritic cell biology and its role in immune-mediated diseases. Clin Transl Immunology, 2020. 9(5): p. e1139.

2. Barrat, F.J. and L. Su, A pathogenic role of plasmacytoid dendritic cells in autoimmunity and chronic viral infection. J Exp Med, 2019. 216(9): p. 1974–1985.

3. Zhou, B., T. Lawrence, and Y. Liang, The Role of Plasmacytoid Dendritic Cells in Cancers. Front Immunol, 2021. 12: p. 749190.

4. Xia, C., et al., Blastic plasmacytoid dendritic cell neoplasm: ontogeny, immunophenotype, genetics and epigenetics, immune dysregulation, skin tropism and therapeutic implications. Leukemia, 2026.

5. Vasilakou, A., et al., The plasmacytoid dendritic cell paradox in cancer: impaired type I interferon responses in a nucleic acid-rich tumor microenvironment. Front Immunol, 2026. 17: p. 1847012.

6. Swarup, N., et al., Cell-Free DNA: Features and Attributes Shaping the Next Frontier in Liquid Biopsy. Mol Diagn Ther, 2025. 29(3): p. 277–290.

7. de Miranda, F.S., et al., Properties and Application of Cell-Free DNA as a Clinical Biomarker. Int J Mol Sci, 2021. 22(17).

8. Malki, Y., et al., The comings and goings of cell-free DNA: Biological and clinical implications. Med, 2026. 7(2): p. 100926.

9. Torfs, K., et al., Neutrophils as critical orchestrators of chronic inflammation. Cell Mol Immunol, 2026. 23(2): p. 123–149.

10. Kahlenberg, J.M. and M.J. Kaplan, Little peptide, big effects: the role of LL-37 in inflammation and autoimmune disease. J Immunol, 2013. 191(10): p. 4895–901.

11. Keshri, A.K., et al., LL-37, the master antimicrobial peptide, its multifaceted role from combating infections to cancer immunity. Int J Antimicrob Agents, 2025. 65(1): p. 107398.

12. Vorobjeva, N.V. and B.V. Chernyak, NETosis: Molecular Mechanisms, Role in Physiology and Pathology. Biochemistry (Mosc), 2020. 85(10): p. 1178–1190.

13. Lande, R., et al., Plasmacytoid dendritic cells sense self-DNA coupled with antimicrobial peptide. Nature, 2007. 449(7162): p. 564–9.

14. Badal, D., et al., Role of DNA-LL37 complexes in the activation of plasmacytoid dendritic cells and monocytes in subjects with type 1 diabetes. Sci Rep, 2020. 10(1): p. 8896.

15. Chan, L.K., et al., Extrachromosomal circular DNA promotes inflammation and hepatocellular carcinoma development. Sci Adv, 2025. 11(42): p. eadw0272.

16. Luo, J., et al., Extrachromosomal circular DNA in cancer drug resistance and its potential clinical implications. Front Oncol, 2022. 12: p. 1092705.

17. Hariprakash, J.M., et al., Functional role of small extrachromosomal circular DNA in colorectal cancer. Proc Natl Acad Sci U S A, 2026. 123(14): p. e2523047123.

18. Petito, V., et al., Inflammatory bowel disease (IBD) is associated with increased intestinal extrachromosomal circular DNA: an emerging biomarker for IBD type and activity. J Crohns Colitis, 2026. 20(2).

19. Ain, Q., et al., Extrachromosomal Circular DNA: Current Knowledge and Implications for CNS Aging and Neurodegeneration. Int J Mol Sci, 2020. 21(7).

20. Wang, Y., et al., eccDNAs are apoptotic products with high innate immunostimulatory activity. Nature, 2021. 599(7884): p. 308–314.

21. Greitens, C., et al., The intracellular journey of DNA delivered with the protein-based transfection system TFAMoplex and Lipofectamine. Int J Pharm, 2025. 681: p. 125855.

22. Dos Santos, C.R., et al., Variation of extrachromosomal circular DNA in cancer cell lines. Comput Struct Biotechnol J, 2023. 21: p. 4207–4214.

23. Li, F., et al., Minicircle HBV cccDNA with a Gaussia luciferase reporter for investigating HBV cccDNA biology and developing cccDNA-targeting drugs. Sci Rep, 2016. 6: p. 36483.

24. Du, Q., A. Kotlyar, and A. Vologodskii, Kinking the double helix by bending deformation. Nucleic Acids Res, 2008. 36(4): p. 1120–8.

25. Carmona-Saez, P., et al., Metagene projection characterizes GEN2.2 and CAL-1 as relevant human plasmacytoid dendritic cell models. Bioinformatics, 2017. 33(23): p. 3691–3695.

26. Kamat, S. and M. Kumari, Repurposing Chloroquine Against Multiple Diseases With Special Attention to SARS-CoV-2 and Associated Toxicity. Front Pharmacol, 2021. 12: p. 576093.

27. Barrat, F.J., et al., Nucleic acids of mammalian origin can act as endogenous ligands for Toll-like receptors and may promote systemic lupus erythematosus. J Exp Med, 2005. 202(8): p. 1131–9.

28. Ma, C., et al., Toll-Like Receptor 9 Inactivation Alleviated Atherosclerotic Progression and Inhibited Macrophage Polarized to M1 Phenotype in ApoE-/- Mice. Dis Markers, 2015. 2015: p. 909572.

29. Schmidt, N.W., et al., Liquid-crystalline ordering of antimicrobial peptide-DNA complexes controls TLR9 activation. Nat Mater, 2015. 14(7): p. 696–700.

30. Swiecki, M. and M. Colonna, The multifaceted biology of plasmacytoid dendritic cells. Nat Rev Immunol, 2015. 15(8): p. 471–85.

31. Bao, M. and Y.J. Liu, Regulation of TLR7/9 signaling in plasmacytoid dendritic cells. Protein Cell, 2013. 4(1): p. 40–52.

32. Combes, A., et al., BAD-LAMP controls TLR9 trafficking and signalling in human plasmacytoid dendritic cells. Nat Commun, 2017. 8(1): p. 913.

33. Doring, Y., et al., Auto-antigenic protein-DNA complexes stimulate plasmacytoid dendritic cells to promote atherosclerosis. Circulation, 2012. 125(13): p. 1673–83.

34. Gilliet, M. and R. Lande, Antimicrobial peptides and self-DNA in autoimmune skin inflammation. Curr Opin Immunol, 2008. 20(4): p. 401–7.

35. Lande, R., et al., Cationic antimicrobial peptides in psoriatic skin cooperate to break innate tolerance to self-DNA. Eur J Immunol, 2015. 45(1): p. 203–13.

36. Lande, R., et al., Neutrophils activate plasmacytoid dendritic cells by releasing self-DNA-peptide complexes in systemic lupus erythematosus. Sci Transl Med, 2011. 3(73): p. 73ra19.

37. Xian, H., et al., Author Correction: Mitochondrial DNA oxidation propagates autoimmunity by enabling plasmacytoid dendritic cells to induce T(FH) differentiation. Nat Immunol, 2026.

38. Lee, E.Y., et al., Helical antimicrobial peptides assemble into protofibril scaffolds that present ordered dsDNA to TLR9. Nat Commun, 2019. 10(1): p. 1012.

39. Leon, S.A., et al., Free DNA in the serum of cancer patients and the effect of therapy. Cancer Res, 1977. 37(3): p. 646–50.

40. Barbany, G., et al., Cell-free tumour DNA testing for early detection of cancer - a potential future tool. J Intern Med, 2019. 286(2): p. 118–136.

41. Gu, Y., et al., The biogenesis of extrachromosomal circular DNA is associated with enhancer-promoter dynamics and chromatin architecture. Int J Biol Macromol, 2026. 364: p. 152307.

42. Yang, L., et al., Extrachromosomal circular DNA: biogenesis, structure, functions and diseases. Signal Transduct Target Ther, 2022. 7(1): p. 342.

43. Hung, K.L., Oncogenes out of context: Cancer genes break free from the regulatory constraints of chromosomes. Science, 2025. 390(6774): p. 686.

44. Luo, Y., et al., Pathological role of excessive DNA as a trigger of keratinocyte proliferation in psoriasis. Clin Exp Immunol, 2020. 202(1): p. 1–10.

45. Peng, Y., et al., Exploring the Role of Extrachromosomal Circular DNA in Human Diseases. Cytogenet Genome Res, 2024. 164(5-6): p. 181–193.

46. Moreno-Angarita, A., C.C. Aragon, and G.J. Tobon, Cathelicidin LL-37: A new important molecule in the pathophysiology of systemic lupus erythematosus. J Transl Autoimmun, 2020. 3: p. 100029.

47. Li, Y., I.C. Berke, and Y. Modis, DNA binding to proteolytically activated TLR9 is sequence-independent and enhanced by DNA curvature. EMBO J, 2012. 31(4): p. 919–31.

48. Kindrachuk, J., et al., Nucleic acids exert a sequence-independent cooperative effect on sequence-dependent activation of Toll-like receptor 9. J Biol Chem, 2007. 282(19): p. 13944–53.

49. Wang, X., et al., Extrachromosomal circular DNA drives ferroptosis and immunity in trophoblasts via the STING pathway in preeclampsia. Biochem Biophys Res Commun, 2025. 781: p. 152522.

50. Wu, Z., et al., Erratum for the Clinical and Translational Medicine “Multi-omics integration reveals the oncogenic role of eccDNAs in diffuse large B-cell lymphoma through STING signalling” by Zijuan Wu et al. Clin Transl Med, 2025. 15(4): p. e70301.

51. Wang, Y., M. Wang, and Y. Zhang, Purification, full-length sequencing and genomic origin mapping of eccDNA. Nat Protoc, 2023. 18(3): p. 683–699.

52. Pazgier, M., et al., Structural and functional analysis of the pro-domain of human cathelicidin, LL-37. Biochemistry, 2013. 52(9): p. 1547–58.

